# A feature-based network analysis and fMRI meta-analysis reveal three distinct types of prosocial decisions

**DOI:** 10.1101/2020.12.09.415034

**Authors:** Shawn A. Rhoads, Jo Cutler, Abigail A. Marsh

## Abstract

Tasks that measure correlates of prosocial decision-making share one common feature: agents can make choices that increase the welfare of a beneficiary. However, prosocial decisions vary widely as a function of other task features. The diverse ways that prosociality is defined and the heterogeneity of prosocial decisions have created challenges for interpreting findings across studies and identifying their neural correlates. To overcome these challenges, we aimed to organize the prosocial decision-making task-space of neuroimaging studies. We conducted a systematic search for studies in which participants made decisions to increase the welfare of others during fMRI. We identified shared and distinct features of these tasks and employed an unsupervised graph-based approach to assess how various forms of prosocial decision-making are related in terms of their low-level components (e.g., task features like potential cost to the agent or potential for reciprocity). Analyses uncovered three clusters of prosocial decisions, which we labeled cooperation, equity, and altruism. This feature-based representation of the task structure was supported by results of a neuroimaging meta-analysis that each type of prosocial decisions recruited diverging neural systems. Results clarify some of the existing heterogeneity in how prosociality is conceptualized and generate insight for future research and task paradigm development.

## Introduction

Prosocial decisions—choices that increase the welfare of others—are universal across cultures (Henrich et al., 2005) and are integral for supporting interpersonal relationships at multiple scales, including between dyads (Declerck et al., 2013; Rusbult & Van Lange, 2003), among groups and social networks (de Waal, 2008; Fehr et al., 2002; Fehr & Camerer, 2007; Fehr & Fischbacher, 2003; FeldmanHall, 2017; Fowler & Christakis, 2010), and within societies (Nowak, 2006). And, while largely conserved across species (Burkart et al., 2014; de Waal, 2008; Hare, 2017), the prevalence and variety of prosociality exhibited by humans is unique (Fehr & Schurtenberger, 2018; Zaki & Mitchell, 2013). Although cognitive and neural processes underlying various forms of prosociality have been studied extensively across disciplines spanning psychology, neuroscience, economics, and biology, the heterogeneity of prosocial decisions has led to inconsistencies in how they are operationalized and categorized (Batson & Powell, 2003; de Waal, 2008; Declerck et al., 2013; Fehr et al., 2002; Fehr & Schmidt, 1999; Marsh, 2016; Parnamets et al., 2020; Rand & Nowak, 2013; Rilling et al., 2002; Ruff & Fehr, 2014; Tricomi & Sullivan-Toole, 2015). This can create challenges when interpreting findings across neuroimaging studies or when attempting to understand how different types of prosocial decisions vary in terms of their underlying processes.

Derived from the Latin stem *pro* and root *socius*, signifying “for a companion”, prosocial decision-making refers to “decisions made for the benefit of another” (*The American Heritage Dictionary of the English Language*, 2000). Laboratory tasks that measure correlates of prosocial decision-making share one common feature: allowing deciding participants (or agents) to make choices that increase the welfare of a beneficiary. However, prosocial decisions vary widely as a function of other task features. For example, although choosing to forgo resources (usually money) to alleviate the suffering of a stranger (in a charitable donation task) versus choosing to contribute money to maximize equity among known members in a group (in a Public Goods Game) both share a common prosocial core of increasing the welfare of others, these decisions diverge along multiple other characteristics. In the first example, a prosocial agent sacrifices resources in response to another person’s distress with the understanding that they will not receive anything in return—suggesting a likely role for empathic concern and planning for prosocial action without any anticipation of reward. In the second example, all prosocial agents are relying on the decisions of others and are hoping to increase the total pool of resources for everyone involved. This suggests a role for monitoring the expected actions of others’ decisions and includes the anticipation of self-rewarding outcomes.

In addition to charitable donation tasks and Public Goods Games, other common prosocial paradigms include Dictator Games, Prisoner’s Dilemmas, and Trust Games. Such tasks can be implemented with multiple variations, and the vast number of combinations of task features is a major source of heterogeneity. This heterogeneity raises the question of whether common mechanisms underlie all prosocial choices. One possibility is that prosocial decisions in each distinct task are supported by distinct mechanisms. But it is more likely that taxonomic clusters exist within the task-space of prosocial decision-making that reflect common underlying neural processes (Cutler & Campbell-Meiklejohn, 2019). One way to identify such clusters would be via a bottom-up approach aimed at characterizing the task structure of prosocial decision-making by analyzing the way specific tasks cluster according to their low-level features. In other words, developing one level of a formal representation (or ontology) of cognitive tasks and their inter-relationships (Poldrack & Yarkoni, 2016; Turner & Laird, 2012). The first goal of this paper was to clarify how different prosocial tasks are inter-related and how their low-level features give rise to broad categories of prosocial decisions. Then, using this information across various studies, we employed an unsupervised graph-based approach to generate a preliminary characterization of the neuroimaging task-space comprised of the distinct and shared task features of prosocial decision-making paradigms. Finally, we conducted an fMRI meta-analysis to identify patterns of distinct and overlapping neural activation that correspond to the identified clusters of prosocial processes.

Breaking prosocial decision processes down into their relevant task features may allow a better understanding of how prosocial decisions are inter-related, and how they diverge. In general, the features that distinguish these tasks involve those related to the *beneficiary* (Is the beneficiary a real or imaginary person (or persons) or an organization like a charity? Is their identity apparent to the agent? Is their need or distress known to the agent?), to the *interaction* (Does the beneficiary also make decisions that will affect ultimate outcome? Will the agent and beneficiary interact only once or more than once?), and to the *outcomes* of the agent’s decision (What is the magnitude of the benefit to the beneficiary? Will the decision result in rewarding outcomes for the agent? Will it be costly? Will the decision conform to social norms, such as equity? How certain is the outcome?). Multiple combinations of these features likely shape the context, motivations, and outcomes of prosocial decisions, and thus should recruit diverging neural systems.

### Features related to the beneficiary

Various features related to the beneficiary of a prosocial decision are known to influence such decisions. Beneficiaries can include specific people, such as close or familiar others (Fareri et al., 2015; Hill et al., 2017; Schreuders et al., 2018; Sharp et al., 2011; Telzer et al., 2011) or in-group members (Balliet et al., 2014; Hackel et al., 2017; Telzer et al., 2015; Wills, Hackel, & Van Bavel, 2018), or can be hypothetical or even non-human (e.g., computers) (Delgado et al., 2005; Fareri et al., 2012). Across contexts, agents are typically more willing to help people than computers (Fareri et al., 2015), and are more willing to help people close to them than strangers (Jones & Rachlin, 2006, 2009; Safin et al., 2013; Strombach et al., 2015). Neural activation during decisions that affect real versus imaginary beneficiaries (e.g., computer) is increased in regions important for theory of mind or inferring the mental states of others such as the temporoparietal junction (TPJ) (FeldmanHall et al., 2012).

In tasks that include real beneficiaries who are previously unknown to the agent, the beneficiary may be another participant in study (Weiland et al., 2012), or an anonymous stranger (Bault et al., 2014; Hutcherson et al., 2015; Strombach et al., 2015) who the agent may have briefly met before the task (Abe et al., 2019; Shaw et al., 2018) or seen in a photograph (Genevsky et al., 2013; Park et al., 2017). Receiving any identifying information about a beneficiary generally increases prosociality, in line with the identifiable victim effect (Jenni & Loewenstein, 1997; Kogut & Ritov, 2005; Lee & Feeley, 2016). This effect also results in greater prosociality toward single individuals versus collectives (Kogut & Ritov, 2005; Lee & Feeley, 2016), including charitable organizations, whether predetermined (Greening et al., 2014), of the agent’s choosing (Kuss et al., 2013), or from a list of charities (Hare et al., 2010; Izuma et al., 2010; Tusche et al., 2016). Increases in prosocial decision-making are particularly robust when the need or distress of the beneficiary is salient (FeldmanHall et al., 2015; Genevsky et al., 2013; Kuss et al., 2015; Tusche et al., 2016). Cues that signal need or distress typically elicit empathic concern, which motivates the desire to alleviate it (Batson, 2011; de Waal, 2008; Marsh, 2016; Preston & de Waal, 2002). This form of empathy is supported by activity in neural regions including the anterior insula, ACC, and pre-supplementary motor area (pre-SMA) (Fallon et al., 2020; Jauniaux et al., 2019; Kogler et al., 2020; Lamm et al., 2011; Schurz et al., 2020) and empathic neural responding predicts prosocial decision-making both in and out of the laboratory (Tusche et al., 2016; Vekaria et al., 2020).

### Features related to the interaction

Aspects of the interaction between agents and beneficiaries (or other agents) in prosocial tasks also influence agents’ decisions, particularly when agents can learn about those with whom they are interacting. In some interactions, only one agent can influence the outcome. For example, in Dictator Games, agents unilaterally allocate resources between themselves and a beneficiary (Engel, 2011). In others, multiple agents can shape the outcome. For example, in social dilemmas or Trust Games, agents can choose to cooperate with others in order to increase the total pool of available resources for everyone involved, or can defect to obtain better outcomes for self (Balliet et al., 2011). Alternatively, in ultimatum games, agents receive feedback about their decisions from beneficiaries, who can accept or reject the offer (Güth et al., 1982).

Some prosocial decisions involve repeated interactions, which, unlike one-shot interactions, provide opportunities to reciprocate or respond to feedback about prior choices (Thielmann et al., 2020). When repeated interactions are expected, it typically motivates cooperation, with agents motivated to pay short-term cooperation “costs” to increase future reciprocity from a partner (Milinski et al., 2001; Rand & Nowak, 2013) and more willing to cooperate with partners who have cooperated previously (Fehr & Schurtenberger, 2018). This may be related to the ability to update expectations of others’ likely behavior, a type of social learning is supported by the subgenual anterior cingulate cortex (ACC) (Christopoulos & King-Casas, 2014). During these repeated interactions, agents may also act to influence interaction partners to reciprocate, for example, when a cooperative exchange is broken and partners coax others back into cooperation via generosity (Bendor et al., 1921; King-Casas et al., 2008).

### Features related to outcomes

Across interaction types, prosocial decisions are also shaped by their anticipated outcomes. In some cases, prosocial decisions may benefit the agent directly. In Prisoner’s Dilemmas or Public Goods Games, for example, decisions to cooperate increase the probability of future reciprocity. In such tasks, agents also take on the role of beneficiaries (Chaudhuri, 2011; Rand & Nowak, 2013), and thus must arbitrate between their own and others’ rewards. These tasks often recruit neural systems that support subjective valuation and reward expectancy, such as the ventromedial PFC and ventral striatum (Parnamets et al., 2020; Wills, Hackel, & Van Bavel, 2018; Wills, Hackel, FeldmanHall, et al., 2018). These tasks also carry an element of uncertainty (Bellucci et al., 2017), with the agent’s outcome often dependent on a beneficiary or trustee’s choices (Mayer et al., 1995). Uncertainty during these decisions may be reflected through activation in the dorsal ACC (Aimone et al., 2014).

Prosocial choices may also yield more abstract rewards, such as conformity to desirable social norms like maximizing equity among multiple parties (via, for example, a 50-50 split of resources) (Fehr & Schurtenberger, 2018; Krupka & Weber, 2013; López-Pérez, 2008). In some cases, agents may choose to act prosocially and forgo resources to avoid deviating from desirable norm, which is known as disadvantageous inequity aversion (Tricomi & Sullivan-Toole, 2015). Equitable interpersonal decisions are thought to engage neural structures involved in computing subjective value such as the medial PFC and ventral striatum, and thus may be motivated through increased intrinsic value placed on the decision (Zaki & Mitchell, 2011), perhaps via their goal of producing increased subjective happiness for agents and beneficiaries (Tabibnia et al., 2007; Tabibnia & Lieberman, 2007). In tasks with repeated interactions, these decisions may also reflect the maintenance of abstract, norm-based rules regarding fairness or reciprocity, in that repeated interactions can enable norms to become established among the interacting parties and maintained in dorsolateral PFC (Guroglu et al., 2014; van den Bos et al., 2009).

In other prosocial decision-making tasks (such as Dictator Games or charitable giving tasks) agents can forgo resources (including money, time, effort, or safety) solely to benefit others. In this case, prosocial choices are made despite certain concrete costs to the agent, often to alleviate the beneficiary’s distress or need. As described above, such decisions are thought to be driven by activation in regions like anterior insula, which represent negative affective states (e.g., pain or distress) of the beneficiary (FeldmanHall et al., 2015; Tusche et al., 2016). Such choices also may yield indirect gains, including increases in mood or well-being (Aknin et al., 2012; Curry et al., 2018; Dunn et al., 2008), possibly related to the vicarious reward of improving the beneficiary’s welfare (Mobbs et al., 2009); such vicarious reward may be supported by activity in ventral striatum and ventromedial PFC.

Given the diversity of extant prosocial decision tasks, two recent meta-analytic studies have been very valuable in describing the neural correlates of prosocial behaviors aggregated across tasks that reflect divergent constellations of the above variables. Belluci and colleagues (2020) aggregated across a wide range of tasks in which participants made decisions about others, rated others’ traits, or judged others’ behaviors in an effort to find neural activation overlap among prosociality, empathy, and mentalizing. They found four regions to be preferentially engaged across the tasks they incorporated: dorsolateral PFC, ventromedial PFC, dorsal posterior cingulate cortex (PCC), and middle cingulate cortex (MCC). Of these regions, they found a conjunction in dorsal PCC activation during tasks involving prosocial behavior and tasks involving mentalizing (understanding another person’s needs and inferring goals across contexts); they also found a conjunction in MCC activation across tasks involving prosocial behavior and tasks involving empathy (resonating with another’s needs) (Bellucci et al., 2020). Activation during prosocial behavior in the dorsolateral PFC and ventromedial PFC did not overlap with activation during mentalizing or empathy tasks. This work identified common neural patterns underlying a range of behaviors related to prosociality, but by not considering key differences among types of prosocial decisions, it was not able to identify whether they are supported by distinct processes. Cutler and Campbell-Meiklejohn (2019) provided preliminary evidence that distinct neural regions do indeed support different forms of prosocial decision-making, finding diverging patterns of activation for prosocial behaviors that do not provide an opportunity to gain extrinsic rewards (and thus likely are intrinsically motivated) versus those with the probability of gaining an extrinsic reward. For example, extrinsically motivated decisions recruited greater activity in striatal regions relative to intrinsically motivated decisions. In contrast, intrinsically motivated decisions recruited increased activation in ventromedial PFC relative to extrinsically motivated decisions. Activation in ventromedial PFC also differentiated these types along a posterior (intrinsic) to anterior (extrinsic) axis.

However, the distinction between extrinsic and intrinsic motivation was determined in advance, rather than being driven by objective features of the data. This is also only one of many possible distinctions among forms of prosocial behavior. An alternative means of investigating neural substrates of various prosocial decision tasks could instead take a more bottom-up approach that identifies distinct clusters of tasks that emerge from statistical variation in their objective features or outcomes. For example, a recent behavioral study analyzed the behavioral outcomes of different economic prosocial tasks (such as the percentage of prosocial decisions during each task, the ratio of other-regarding to self-regarding decisions in each, average monetary donations, or summary scores of self-reported measures). Using factor analysis, they determined that the prosocial tasks clustered into four factors that the authors termed: altruistically motivated prosocial behavior, norm-motivated prosocial behavior, strategically motivated prosocial behavior, and self-reported prosocial behavior (Böckler et al., 2016).

We sought to use a similar bottom-up approach to meta-analytically investigate the neural correlates of prosocial decision-making during fMRI. We focused on objective features that distinguish the tasks themselves, which included features related to outcomes of decisions, to the beneficiaries of the decision, and to the interaction between agents and beneficiaries. We first compiled data from 43 unique fMRI studies of prosocial decision-making (including 25 maps and 18 coordinate tables across 1,423 participants). We then dummy-coded task features related to the beneficiary, interaction, and outcome of each decision and employed a data-driven, graph-based approach to identify clusters of studies based on their overlapping versus distinct task features. As described in detail below, this approach indicated that the prosocial decision-making task-space comprises three clusters, which we labeled cooperation, equity, and altruism. We next used a meta-analytic approach that combined group-level statistical parametric images with reported peak-coordinates to identify divergent neural activation patterns across these clusters of studies. In so doing, the present study resolves some discrepancies in how prosocial decisions are conceptualized, expands understanding regarding how prosocial decisions are related and distinct, and generates insight for future research.

## Method

### Literature search and study selection

A literature search using PubMed identified research published prior to June 2019 using keywords either (“fMRI” or “neur*”) and one of the following: (“prosocial”, “altruis*”, “trust game”, “fairness”, “reciproc*”, “cooperat*”, “charitable”, “public goods”, “dictator”, “ultimatum”, “prisoner*”)^†^. The search returned 201 articles. We removed 124 articles that did not meet key criteria, such as non-neuroimaging studies, neuroimaging studies that did not use functional magnetic resonance imaging (fMRI), and literature reviews or meta-analyses. Independently, we identified 123 potential articles from the reference lists of the remaining 77 articles. After removing all duplicate titles from the combined lists, 146 articles remained. We then selected only articles that reported novel whole-brain fMRI data (i.e., data only published once) that were collected while participants made decisions that benefitted another individual (prosocial decisions). We also limited our search to include only data from studies that were able to examine differences in activation during prosocial decisions relative to decisions that benefited the agent alone (selfish decisions). In some cases, this was a contrast between prosocial choices and selfish choices within a task condition or parametric modulation of the amount given. For other studies, the contrast was between decisions during a prosocial condition and a self-only condition. We did not include contrasts involving alternate control conditions (e.g., rest, visuomotor controls), even when these were available, due to significant variation in brain activation (Cutler & Campbell-Meiklejohn, 2019). Upon review of the remaining 146 articles, 69 were identified that met our inclusion criteria.

We sent emails to the corresponding authors of all included studies to request unthresholded, group-level t-statistic map(s) from the study that best fit our criteria. For studies that included pharmacological manipulations or clinical populations, we requested data from only the control group. If maps were not available, we requested coordinates for contrasts of interests or extracted them from manuscripts. If a coordinate table reported Z-scores or Talaraich coordinates, peak values were transformed to *t*-statistics and MNI coordinates, respectively. If the contrast of interest was reported in both directions (e.g. cooperate > defect and defect > cooperate), the selfish contrast peaks were assigned as negative t-values. Ultimately, we obtained the necessary data from 43 unique fMRI studies, including 25 maps and 18 coordinate tables that included data from 1,423 subjects (Table 1)

**Table 1.** Descriptions of studies included in meta-analysis

| Study | N | Proportion female | Map or peak | Task | T | FWHM | Program | Sig | Contrast selected | Cluster Label |
| --- | --- | --- | --- | --- | --- | --- | --- | --- | --- | --- |
| (Abe et al., 2019) | 19 | 9 / 19 | map | Joint force-production task | 3T | 8mm<sup>3</sup> | SPM12 | Pcorr < 0.001 | Joint performance vs single performance | C |
| (Aimone et al., 2014) | 28 | 15 / 30 total | peak | Trust Game, one-shot, binary (investor) | 3T | 8mm<sup>3</sup> | SPM8 | Puncorr < .0001 | Share vs keep | C |
| (Bault et al., 2014) | 25 | 12 / 29 total | map | Public Goods Game | 3T | 5mm<sup>3</sup> | FSL | Pcorr < 0.05 (FWE) | PM monetary choice | C |
| (Chen et al., 2016) | 93 | 50 / 104 total | map | Prisoner's Dilemma, iterative | 3T | 5mm<sup>3</sup> | FSL | Pcorr < 0.05 (FWE) | Cooperation vs defect (placebo only) | C |
| (Decety et al., 2004) | 12 | 6 / 12 | peak | Cooperative Pattern Game | 3T | 4.5 x 4.5 x 6 mm | SPM2 | Pcorr < 0.05 | Cooperation vs competition | C |
| (Delgado et al., 2005) | 12 | 6 / 14 total | peak | Trust Game, binary, iterative (investor) | 3T | 4mm<sup>3</sup> | Brain Voyager | Puncorr < 0.001 | Share vs keep | C |
| (Fareri et al., 2015) | 26 | 14 / 26 | map | Trust Game, binary, iterative (investor) | 3T | 4mm<sup>3</sup> | Brain Voyager | N/A | Share vs keep (across all partners) | C |
| (FeldmanHall et al., 2012) | 14 | 8 / 14 | peak | Your Pain, My Gain Task | 3T | 8mm<sup>3</sup> | SPM | Pcorr < 0.05 | PM monetary choices, no covariates | A |
| (FeldmanHall et al., 2015) | 17 | 11 / 17 | peak | Your Pain, My Gain Task | 3T | 8mm<sup>3</sup> | SPM5 | Pcorr < 0.05 (FWE) | PM monetary choice | A |
| (Fermin et al., 2016) | 33 | 18 / 33 | map | Prisoner's Dilemma, one-shot | 3T | 8mm<sup>3</sup> | SPM8 | Puncorr < .005 | Cooperation vs defect | C |
| (Fouragnan et al., 2013) | 18 | 0 / 18 | peak | Trust Game (with priors manipulation) | 4T | 8mm<sup>3</sup> | SPM8 | Pcorr < 0.005 | Share vs keep | C |
| (Garbarini et al., 2014) | 16 | 8 / 16 | map | Trust Game, one-shot (responder) | 1.5T | 4mm<sup>3</sup> | Brain Voyager | Pcorr < 0.05 | Reciprocate vs defect | E |
| (Genevsky et al., 2013) | 11 | 6 / 11 | peak | Charitable donation task with identifiable victim | 3T | 4mm<sup>3</sup> | AFNI | Puncorr < 0.005 | Donation vs no donation | A |
| (Greening et al., 2014) | 18 | 9 / 18 | peak | Charity task | 3T | 4mm<sup>3</sup> | AFNI | Pcorr < 0.05 | Charity vs self | A |
| (Guroglu et al., 2014) | 22 | 17 / 28 total | peak | Modified Dictator Game (self-maximizing inequity (SMI) game) | 3T | 8mm<sup>3</sup> | SPM8 | Puncorr < 0.005 | Equal split vs unequal | E |
| (Hare et al., 2010) | 22 | 22 / 22 | map | Charitable donation task | 3T | 8mm<sup>3</sup> | SPM5 | Puncorr < 0.0005 | PM monetary choice | A |
| (Hutcherson et al., 2015) | 51 | 0 / 51 | map | Modified Dictator Game, with probable outcomes | 3T | 8mm^3 | SPM8 | Puncorr < 0.05 | Prosocial choice vs self | A |
| (Izuma et al., 2010) | 23 | 12 / 23 | map | Charitable donation task in presence or absence of observer | 3T | 6mm^3 | SPM5 | Puncorr < 0.001 | Donation vs no donation (no observers) | A |
| (Koban et al., 2014) | 17 | 10 / 22 total | map | Modified Dictator Game, Sharing/keeping during conflict | 3T | 8mm^3 | SPM8 | Puncorr < 0.001 | Equal split vs self during human interpersonal conflict | E |
| (Kuss et al., 2013) | 33 | 14 / 33 | map | Charitable donation task, with probable outcomes | 3T | 6mm^3 | SPM8 | Puncorr < 0.001 | Costly donation vs self | A |
| (Lee et al., 2018) | 16 | 0 / 16 | map | Tetris-like game, with self, helping, and harming conditions | 3T | 8mm^3 | SPM5 | Pcorr < 0.05 | Help vs self | A |
| (Lelieveld et al., 2013) | 26 | 17 / 26 | map | Dictator Game with emotional manipulation | 3T | 6mm^3 | SPM5 | Puncorr < 0.001 | Equal split vs unequal split | E |
| (Morishima et al., 2012) | 27 | 17 / 30 total | map | Dictator Game | 3T | 8mm^3 | SPM8 | Pcorr < 0.05 (FWE) | Donation vs self | A |
| (Park et al., 2017) | 159 | 100 / 166 total | map | Dictator Game with Faces | 3T | 4mm^3 | AFNI | Puncorr < 0.005 | Giving vs not giving (offer only) | A |
| (Ramsøy et al., 2015) | 30 | 14 / 30 | peak | Prisoner's Dilemma, with belief prompts about partner's decision | 3T | 8mm^3 | SPM8 | Puncorr < 0.001 | Cooperation vs defect | C |
| (Schneider-Hassloff et al., 2015) | 164 | 78 / 164 | map | Prisoner's Dilemma with measures of attachment style | 3T | 8mm^3 | SPM8 | Puncorr < 0.001 | Cooperation vs defect | C |
| (Schreuders et al., 2018) | 22 | 12 / 27 total | map | Modified Dictator Game (self-maximizing inequity game) | 3T | 8mm^3 | SPM8 | Pcorr < 0.05 | Equal split vs self (across interaction partners) | E |
| (Schreuders et al., 2019) | 39 | 29 / 50 total | map | Modified Dictator Game (self-maximizing inequity game) | 3T | 8mm^3 | SPM8 | Pcorr < 0.001 | Equal split vs self (across interaction partners) | E |
| (Sharp et al., 2011) | 20 | 0 / 20 | map | Trust Game (investor) | 3T | 4mm^3 | AFNI | Puncorr < 0.001 | Share vs keep (controls) | C |
| (Shaw et al., 2018) | 38 | 0 / 38 | peak | Ultimatum Game (proposer) | 3T | 5mm^3 | FSL (preproc) SPM12 (GLM) | Pcorr < 0.001 (FWE) | PM monetary choice (proposers only) | C |
| (Smith-Collins et al., 2013) | 24 | 24 / 24 | peak | Trust Game, iterative (investor) | 1.5T | 8mm<sup>3</sup> | SPM8 | Puncorr < .001 | Cooperation vs defect | C |
| (Stanley et al., 2012) | 40 | 22 / 40 | map | Trust Game, one-shot (investor) | 3T | 6mm<sup>3</sup> | SPM8 | Pcorr < .05 (FWE) | PM monetary choices (only human partners) | C |
| (Strombach et al., 2015) | 27 | 13 / 27 | map | Modified Dictator Game with social distance manipulation | 3T | 8mm<sup>3</sup> | SPM8 | Pcorr < 0.005 | Equal split vs self | E |
| (Telzer et al., 2011) | 25 | 13 / 25 | map | Family Assistance Task | 3T | 8mm<sup>3</sup> | SPM5 | Pcorr < 0.05 | Costly donation vs self | A |
| (Telzer et al., 2015) | 29 | 13 / 29 | map | Modified Dictator Game with group membership manipulation | 3T | 8mm<sup>3</sup> | SPM8 | Pcorr < 0.05 | Donation vs self | A |
| (Tusche et al., 2016) | 23 | 15 / 33 total | map | Charitable donation task | 3T | 8mm<sup>3</sup> | SPM8 | Pcorr < .05 (FWE) | High donation vs low donation | A |
| (van den Bos et al., 2009) | 18 | 9 / 18 | peak | Trust Game, one-shot (responder) | 3T | 6mm<sup>3</sup> | SPM2 | Puncorr < 0.001 | Reciprocate vs defect | C |
| (Watanabe et al., 2014) | 48 | N/A | peak | Pay-it-forward + reputation-based indirect reciprocity game | 3T | 8mm<sup>3</sup> | SPM8 | Pcorr < 0.05 (FWE) | Cooperation vs defect (pay-it-forward & reputation-based conditions) | C |
| (Weiland et al., 2012) | 14 | 8 / 14 | peak | Dictator Game | 3T | 8mm<sup>3</sup> | Brain Voyager | Puncorr < 0.005 | Fair split (6:6, 7:5) vs unfair (8:4, 9:3, 10:2, 11:1) | E |
| (Will et al., 2016) | 43 | 17 / 43 | map | Dictator Game after Cyberball | 3T | 8mm<sup>3</sup> | SPM8 | Puncorr < .001 | Equitable choice vs inequitable | E |
| (Wills, Hackel, & Van Bavel, 2018) | 42 | 32 / 47 | peak | Public Goods Game | 3T | 6mm<sup>3</sup> | SPM12 | Pcorr < .05 | Give vs keep | C |
| (Wittmann et al., 2016) | 24 | 9 / 24 | peak | Cooperation Task | 3T | 5mm<sup>3</sup> | FSL | Pcorr < 0.05 (FWE) | Cooperation versus competition | C |
| (Zaki & Mitchell, 2011) | 15 | 6 / 15 | peak | Modified Dictator Game | 3T | 6mm<sup>3</sup> | SPM | Pcorr < 0.005 | Equitable choice vs inequitable | E |

**Note.** Some studies only reported gender as a proportion of the total sample, but not the final analytic sample. Researcher labels: C=Cooperative; E=Equitable; A=Altruistic

### Identifying task features across studies

We first reviewed the details of the methodologies of all available tasks and identified 13 distinct task features that varied among existing prosocial decision-making tasks across neuroimaging studies (Figure 1). These task features can be broken down into those vary as a function of the beneficiary, the interaction, and the outcome. Although in theory features related to the agent can also vary, all participants included in the present study were healthy control adults, and we did not identify consistent features related to these participants—for example, consistent individual difference measures—in the available literature. Four independent raters dummy coded (“present” or “absent”) the 13 features during the prosocial decision phase for the available contrast in each study with high initial agreement among coders (*ICC*=.82, CI_95%_=[.79,.84]). Discrepancies in the initial coding were then resolved through a consensus agreement across each of the four coders.

**Figure 1.**
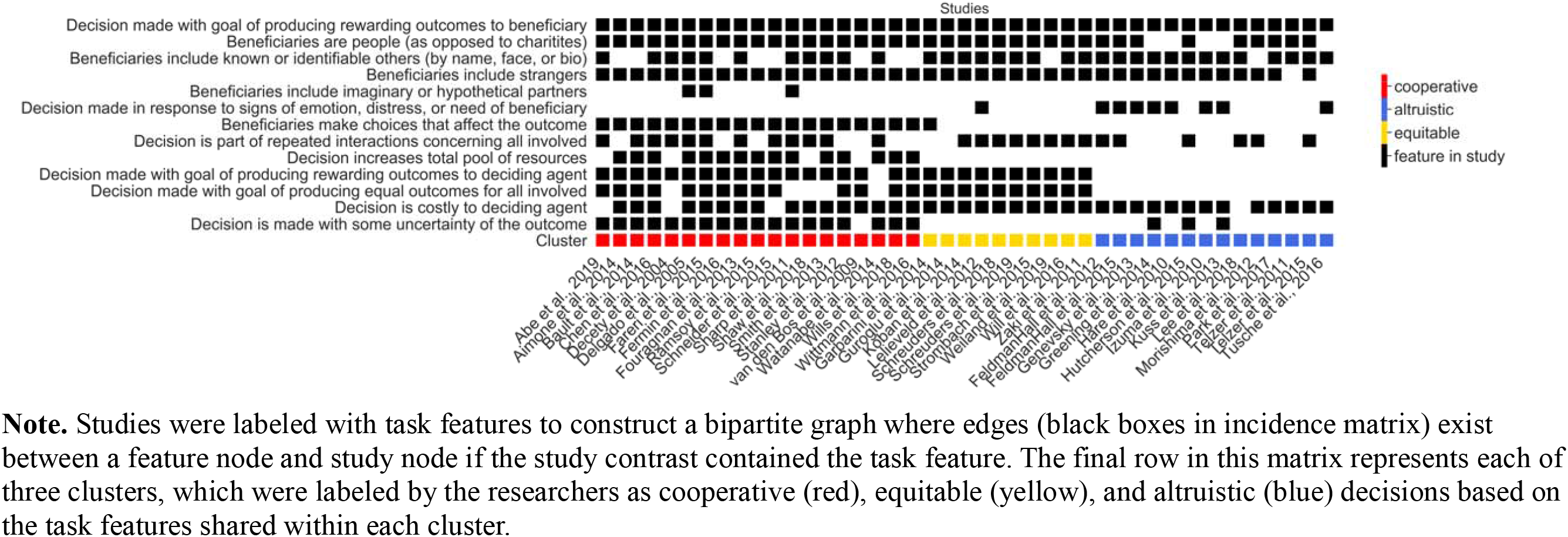
Bipartite feature x study incidence matrix.

### Identifying clusters within the task structure of prosocial decision-making

To assess differential clusters of prosocial decisions based on their task features, we next applied an unsupervised, graph-based approach. Importantly, we selected an approach that allowed us to identify potential clusters of studies among tasks that all shared a common feature of “producing a rewarding outcome to the beneficiary” (a crucial criterion for inclusion in our meta-analysis). Thus, we sought to construct a fully-connected, weighted graph with studies as nodes and the degree of overlapping task features as weighted edges. To accomplish this, we first used the identified features to construct a bipartite graph. This graph contained two sets of nodes: nodes representing the 13 different task features and nodes representing the 43 different prosocial decision study contrasts. In this graph, an edge exists between a feature node and a study node if the study contrast contained the task feature (Figure 1 depicts this graph in matrix form). Next, we projected this bipartite graph onto a weighted network of studies, where edge weights between studies represented the Dice similarity coefficient (Dice, 1945) or the degree of overlapping task features relative to the total possible task features (Figure 2). We then ran the Louvain community detection algorithm (Blondel et al., 2008)—a common unsupervised clustering algorithm used across the biological, psychological, and social sciences that detects clustering of nodes in a fully-connected, weighted graph (Allegra et al., 2020; Gonzalez-Castillo et al., 2019; Ito et al., 2017; Matthews et al., 2016; Miele et al., 2019; Paquola et al., 2019; Tian et al., 2020; Wang et al., 2020). This approach assigned study nodes to clusters in two steps. First, the algorithm finds small clusters of studies by optimizing local modularity. Second, it aggregates studies of the same cluster in a hierarchical fashion and builds a new network whose nodes are these clusters. These steps were repeated iteratively until the global modularity was maximized (i.e., global modularity is achieved when the connections between studies within the same clusters are strongest and connections between nodes in different clusters are weakest). The resulting hierarchy of studies across clusters in the study network is depicted in Figure S1. To assess the stability of the identified three-cluster solution, we implemented a Jackknife sensitivity analysis, which exhaustively left-one-study-out prior to generating the study network and applying Louvain community detection. We then computed (1) the percentage of iterations that yielded the same number of clusters (three clusters) in the study network and (2) the proportion of studies that switched to a different cluster.

**Figure 2.**
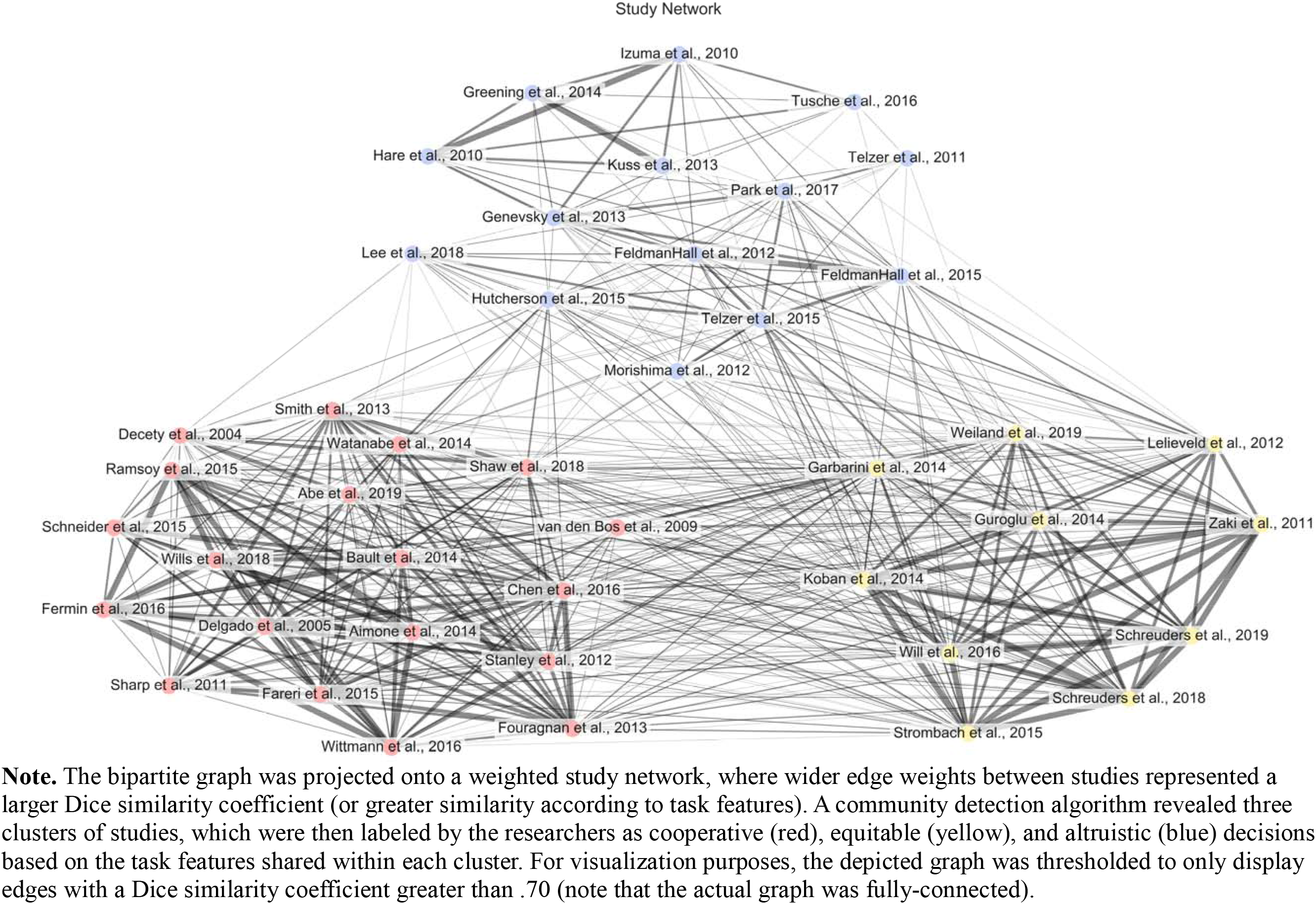
Graph depiction of study network generated from overlapping task features.

### Neuroimaging preprocessing and meta-analyses

We next conducted meta-analyses combining reported peak information (coordinates and *t*-statistics) with original statistical parametric maps using the Anisotropic Effect Size Signed Differential Mapping software (AES-SDM, version 5.141; Radua et al., 2014). We selected this analytical technique rather than alternatives, such as coordinate-based activation likelihood (Eickhoff et al., 2009; Turkeltaub et al., 2002), because this approach enabled the utilization of precise, continuous estimates of effect sizes, assessment of between-study heterogeneity, and identification of potential publication bias (Radua et al., 2012). Using AES-SDM, within-study, voxel-level maps of effect sizes (*Hedge’s g*) and their variances were re-created for each study. When only reported coordinates and statistics were available for a study, we calculated the effect size at each peak and estimated effect sizes in neighboring voxels based on the Euclidean distance between voxels and the peak using a 20mm FHWM Gaussian function (Radua et al., 2012, 2014). This method of estimation is similar to the estimation of activation likelihood used in peak-probability meta-analytic methods, but the use of effect sizes in the calculation increases the accuracy of estimation of the true signal (Radua et al., 2012). When the *t*-statistics of the peak coordinates were unknown (one study: Delgado et al., 2005), we imputed the effect size with the extent threshold reported in the study.

We conducted three random-effects models to compute a meta-analytic activation for each prosocial category identified using the Louvain community detection algorithm. Each individual study was weighted by the inverse sum of its variance plus the between-study variance as obtained by the DerSimonian-Laird estimator of heterogeneity (DerSimonian & Laird, 1986). Within this random-effects framework, studies with larger sample sizes or lower variability contribute more and effects are assumed to randomly vary between study samples. To assess statistical significance, we implemented a modified permutation test that empirically estimated a null distribution for each meta-analytic brain map. We thus tested the hypothesis that each map’s true effect sizes were not the result of a random spatial association among studies within a prosocial category. We applied a threshold of *p*<.005 as recommended by Radua et al. (2012) to optimally balance specificity and sensitivity while yielding results approximately equivalent to *p*<.05 corrected for multiple comparisons. Reported z-scores are specified as SDM-Z, as they do not follow a standard normal distribution. We also conducted three pairwise comparisons of activation maps across each of the three prosocial categories, which followed the same procedures (see Supplemental Table S1).

The effect size maps were imported into AFNI (Cox, 1996) and a conjunction analysis was conducted to examine the overlap of consistently activated regions across altruism, cooperation, and equity. Conjunction was determined using 3dcalc by overlaying the thresholded meta-analytic maps for each category to determine activation overlay.

## Results

### Clustering tasks into categories of prosocial decision-making

The Louvain clustering algorithm revealed three clusters of prosocial decision-making tasks (Figure 2). Upon inspection, we labeled these clusters based on the task features (or lack of features) shared among decisions within each cluster. Decisions in the first cluster (red; N=19; 8 maps, 11 coordinate tables) involved multiple agents acting prosocially to maximize resources. These decisions were characterized by features like: outcomes depended on the decisions of others and decisions in conditions of uncertainty. Tasks included Prisoner’s Dilemmas, Public Goods Games, Ultimatum Games played by proposers, Trust Games, and non-economic cooperative tasks. We labeled this cluster: *cooperative decisions*. A second cluster of decisions (yellow; N=10; 7 maps, 3 coordinate tables) were those in which unilateral decisions to apportion resources equally were possible. These decisions were characterized by features like: adherence to social norms such as producing equitable outcomes for the agent and beneficiary and unilateral decisions made by a single agent (and that thus resulted in no uncertainty). Tasks included Dictator Games in which 50-50 splits were possible. We labeled this cluster: *equitable decisions*. Finally, a third cluster of decisions (N=14; 10 maps, 4 coordinate tables) were those in which agents made unilateral decisions to forgo resources for others. These decisions were characterized by losses without receiving anything in return and decisions that were made in response to the need or distress of the beneficiaries. Tasks included charitable donation tasks, Dictator Games in which 50-50 splits were not possible, Your Pain, My Gain tasks, and assistance tasks. We labeled this cluster: *altruistic decisions*. We will refer to each cluster using these terms, with the recognition that alternate descriptions for each cluster could also be appropriate. For example, the cluster of tasks we labeled “cooperative” could alternately be labeled “strategic” (Böckler et al., 2016). Crucially, sensitivity analyses (iteratively leaving one study out prior to generating the network and applying community detection) revealed that 100% of iterations yielded a three-cluster solution. The stability of these clusters was also high with 97.7% of nodes in the study network remaining in the same cluster; only one study (van den Bos et al., 2009) out of the 43 studies ever switched from the cluster labeled “cooperative” to the cluster labeled “equity”.

### Meta-analyses

#### Neural correlates of cooperative decisions

In the cooperative decision cluster, prosocial decisions (in contrast to selfish decisions) were associated with increased activation in right inferior frontal gyrus, bilateral subgenual ACC, left ventral striatum (including caudate nucleus), bilateral insula, bilateral MCC, left supramarginal gyrus extending to superior temporal gyrus, left lateral postcentral gyrus, bilateral ventral tegmental area (VTA), left thalamus, left precuneus, right cerebellum lobule VIII, and bilateral occipital cortex. We did not find significantly increased activation in any region during selfish versus cooperative decision-making (Figure 3, Table 2).

**Figure 3.**
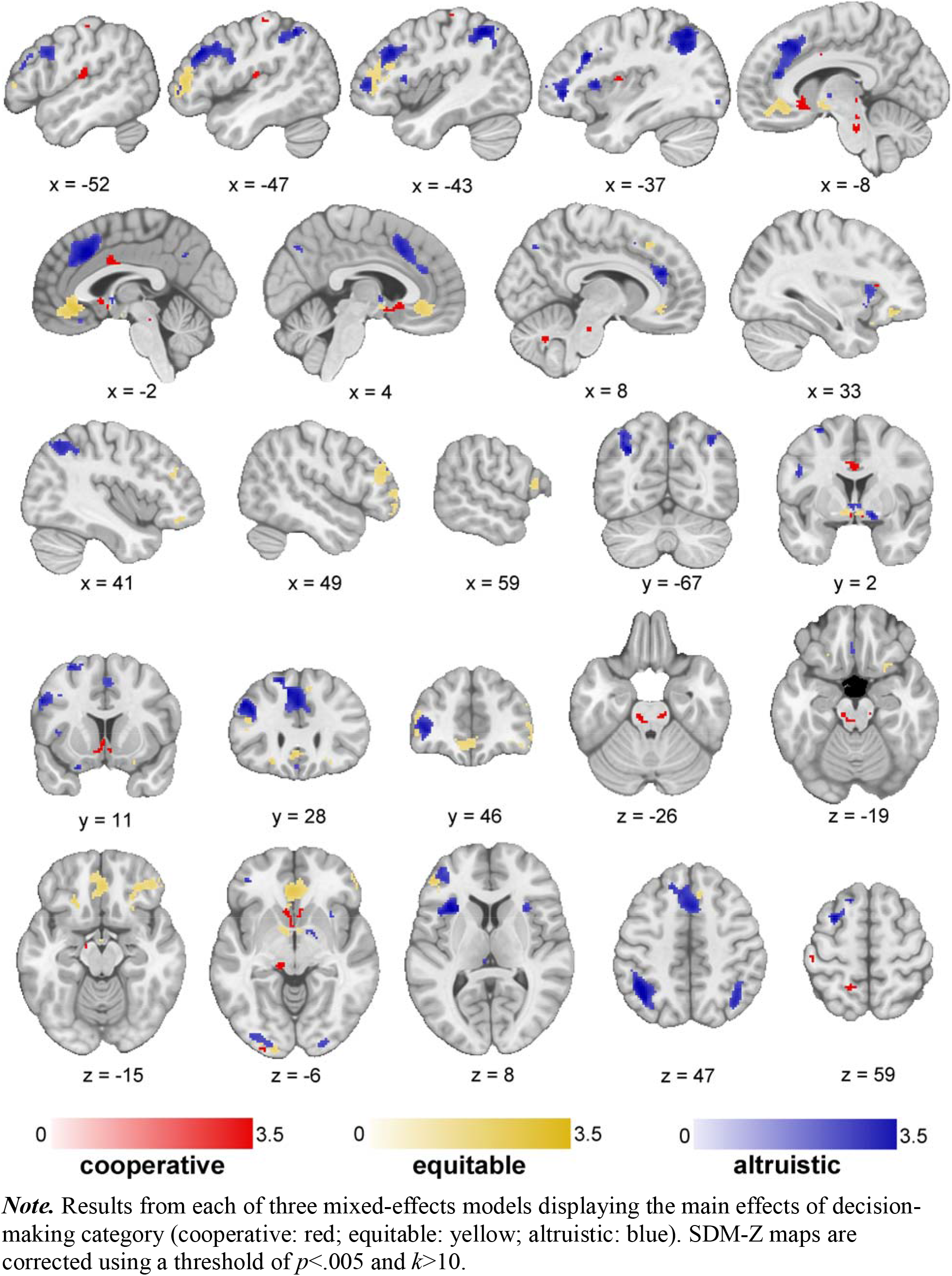
Thresholded results from meta-analyses.

**Table 2.** Results from the meta-analyses.

> selfish decisions (increased activity)
| Region | Brodmann Area | SDM-Z | P | Voxels | MNI-x | MNI-y | MNI-z |
| --- | --- | --- | --- | --- | --- | --- | --- |
| Inferior frontal gyrus, triangular part extending to insula (R) | 48 | 3.824 | 0.000581 | 11 | 34 | 24 | 10 |
| Ventral striatum extending to caudate and subgenual cingulate (L) |  | 4.682 | 0.000021 | 89 | -6 | 16 | -2 |
| Subgenual anterior cingulate cortex extending to olfactory cortex (R) | 25 | 3.65 | 0.001053 | 28 | 4 | 12 | -10 |
| Middle cingulate cortex extending to paracingulate gyri (L, R) | 24 | 3.853 | 0.000523 | 45 | -2 | 4 | 32 |
| Insula (L) | 48 | 3.497 | 0.001740 | 14 | -38 | -4 | 12 |
| Hippocampus (L) |  | 3.408 | 0.002305 | 11 | -16 | -12 | -12 |
| Ventral tegmental area (L) |  | 4.102 | 0.000212 | 51 | -10 | -24 | -28 |
| Ventral tegmental area (R) | 30 | 3.814 | 0.000600 | 30 | 12 | -24 | -22 |
| Postcentral gyrus, lateral part (L) | 3 | 3.861 | 0.000509 | 14 | -50 | -24 | 60 |
| Supramarginal gyrus extending to superior temporal gyrus (L) | 48 | 3.888 | 0.000464 | 95 | -60 | -26 | 22 |
| Thalamus (L) | 50 | 3.931 | 0.000400 | 42 | -14 | -28 | -6 |
| White matter (middle cerebellar peduncles) |  | 3.614 | 0.001188 | 13 | 14 | -30 | -34 |
| Precuneus (L) |  | 3.744 | 0.000764 | 14 | -14 | -50 | 58 |
| Cerebellum lobule VIII (R) |  | 3.568 | 0.001378 | 15 | 8 | -62 | -34 |
| Middle occipital gyrus (L) | 19 | 4.292 | 0.000103 | 49 | -22 | -88 | 18 |
| Middle occipital gyrus (L, R) | 18 | 3.736 | 0.000785 | 11 | -28 | -98 | -6 |

**Equitable > selfish decisions (increased activity)**
| Region | Brodmann Area | SDM-Z | P | Voxels | MNI-x | MNI-y | MNI-z |
| --- | --- | --- | --- | --- | --- | --- | --- |
| Inferior frontal gyrus, triangular part (L) | 45 | 2.366 | 0.000113 | 305 | -48 | 42 | 8 |
| Middle frontal gyrus (R) | 45 | 2.093 | 0.000468 | 205 | 48 | 40 | 20 |
| Middle frontal gyrus extending to superior orbital gyrus (R) | 10, 11 | 2.391 | 0.000098 | 234 | 30 | 38 | -12 |
| Anterior cingulate cortex extending to paracingulate gyri (L, R) | 11 | 2.731 | 0.000014 | 488 | -2 | 32 | -10 |
| Inferior frontal gyrus, orbital part (L) | 11 | 2.302 | 0.000158 | 24 | -24 | 26 | -16 |
| Superior frontal gyrus, medial part (R) | 8 | 1.957 | 0.000904 | 18 | 8 | 26 | 48 |
| Inferior frontal gyrus, opercular part (R) | 44 | 1.996 | 0.000748 | 36 | 60 | 16 | 14 |
| Striatum (L) |  | 1.994 | 0.000752 | 68 | -6 | -6 | -8 |
| Middle occipital gyrus (L) | 18 | 2.39 | 0.000099 | 113 | -18 | -98 | -2 |

**Equitable < selfish decisions (decreased activity)**
| Region | Brodmann Area | SDM-Z | P | Voxels | MNI-x | MNI-y | MNI-z |
| --- | --- | --- | --- | --- | --- | --- | --- |
| Middle frontal gyrus (L) | 46 | -2.534 | 0.000820 | 125 | -22 | 42 | 26 |
| Rolandic operculum (L) |  | -2.341 | 0.001943 | 36 | -52 | 0 | 16 |
| Inferior temporal gyrus (L) | 20 | -2.548 | 0.000771 | 15 | -48 | -2 | -32 |
| Precentral gyrus (R) | 6 | -2.533 | 0.000827 | 32 | 48 | -4 | 40 |
| Supplementary motor area (L) | 6 | -2.831 | 0.000197 | 252 | -6 | -4 | 68 |
| Precentral gyrus (L) | 6 | -2.546 | 0.000778 | 27 | -50 | -8 | 52 |
| Precentral gyrus (L) | 6 | -2.518 | 0.000883 | 17 | -38 | -12 | 40 |
| Caudate nucleus (R) |  | -3.431 | 0.000007 | 91 | 18 | -16 | 24 |
| Precentral gyrus (L) | 6 | -2.355 | 0.001822 | 34 | -30 | -18 | 60 |
| Inferior temporal gyrus (L) | 20 | -2.449 | 0.001206 | 12 | -44 | -18 | -24 |
| Thalamus extending to caudate (L) |  | -3.007 | 0.000079 | 288 | -14 | -20 | 20 |
| Posterior insula (R) |  | -3.078 | 0.000054 | 27 | 32 | -28 | 18 |
| Supramarginal gyrus (R) | 48 | -2.651 | 0.000474 | 70 | 54 | -36 | 28 |
| Posterior cingulate cortex, ventral portion (R) | 30 | -2.347 | 0.001894 | 10 | 14 | -38 | 6 |
| Posterior superior temporal sulcus (R) | 39 | -3.021 | 0.000073 | 65 | 48 | -40 | -2 |
| Supramarginal gyrus (L) | 48 | -2.581 | 0.000663 | 84 | -50 | -40 | 28 |
| Inferior temporal gyrus (L) | 20 | -3.117 | 0.000043 | 105 | -42 | -44 | -10 |
| Posterior superior temporal sulcus (R) | 39 | -2.428 | 0.001329 | 11 | 44 | -48 | 12 |
| Postcentral gyrus (L) | 5 | -2.937 | 0.000113 | 182 | -20 | -52 | 64 |
| Posterior superior temporal sulcus (L) | 39 | -2.592 | 0.000627 | 27 | -40 | -60 | 12 |
| Extrastriate cortex extending to parahippocampal gyrus (R) | 36 | -3.239 | 0.000021 | 211 | 30 | -62 | 6 |
| Primary visual cortex (L) | 17 | -2.488 | 0.00101 | 57 | -24 | -68 | 8 |

**Altruistic > selfish decisions (increased activity)**
| Region | Brodmann Area | SDM-Z | P | Voxels | MNI-x | MNI-y | MNI-z |
| --- | --- | --- | --- | --- | --- | --- | --- |
| Middle orbital gyrus (L) |  | 3.397 | 0.000130 | 200 | -38 | 44 | 0 |
| Middle frontal gyrus (R) | 45 | 2.827 | 0.001091 | 31 | 46 | 40 | 28 |
| Anterior cingulate cortex extending to paracingulate gyri and pre-supplementary motor area (L, R) | 32 | 3.871 | 0.000021 | 1278 | 10 | 36 | 22 |
| Gyrus rectus, ventromedial (L, R) | 11 | 2.564 | 0.002750 | 10 | -2 | 30 | -18 |
| Middle frontal gyrus (L) | 44 | 3.761 | 0.000031 | 568 | -40 | 24 | 32 |
| Anterior insula (L) | 48 | 3.961 | 0.000014 | 203 | -34 | 20 | 8 |
| Anterior insula (R) | 48 | 2.923 | 0.000773 | 64 | 30 | 18 | 6 |
| Inferior frontal gyrus, orbital part (L) | 38 | 2.982 | 0.000624 | 12 | -24 | 12 | -24 |
| Ventral striatum (R) |  | 3.15 | 0.000331 | 50 | 14 | 2 | -8 |
| Thalamus (L) |  | 2.814 | 0.001146 | 40 | -6 | -6 | 0 |
| Thalamus (L) |  | 2.803 | 0.001191 | 23 | -14 | -30 | 16 |
| Inferior parietal (excluding supramarginal and angular) gyri (L) | 40 | 3.917 | 0.000017 | 705 | -40 | -54 | 50 |
| Inferior parietal (excluding supramarginal and angular) gyri (R) | 40 | 3.105 | 0.000394 | 219 | 42 | -56 | 52 |
| Precuneus (L) | 7 | 2.83 | 0.001082 | 13 | -4 | -60 | 38 |
| Precuneus (R) | 7 | 2.96 | 0.000673 | 26 | 6 | -70 | 44 |
| Middle occipital gyrus (L) | 18 | 2.761 | 0.001381 | 194 | -22 | -96 | 0 |
| Inferior occipital gyrus (R) | 18 | 2.594 | 0.002483 | 65 | 24 | -96 | -6 |

**Altruistic < selfish decisions (decreased activity)**
| Region | Brodmann Area | SDM-Z | P | Voxels | MNI-x | MNI-y | MNI-z |
| --- | --- | --- | --- | --- | --- | --- | --- |
| Superior frontal gyrus, dorsolateral (R) | 9 | -2.602 | 0.002379 | 13 | 18 | 50 | 30 |
| Inferior frontal gyrus, pars triangularis part (R) | 48 | -3.033 | 0.000539 | 104 | 48 | 34 | 6 |
| Rolandic operculum (L) | 6, 48 | -3.372 | 0.000147 | 38 | -56 | 0 | 8 |
| Posterior insula (L) | 84 | -2.627 | 0.002198 | 15 | -42 | 0 | 8 |
| Anterior superior temporal sulcus (R) |  | -3.524 | 0.000080 | 69 | 48 | -2 | -14 |
| Putamen (lenticular nucleus) (R) | 48 | -2.541 | 0.002913 | 10 | 30 | -8 | 2 |
| Posterior insula (R) | 48 | -3.902 | 0.000016 | 780 | 42 | -10 | 2 |
| Superior frontal gyrus, dorsolateral (R) | 6 | -3.119 | 0.000394 | 13 | 18 | -10 | 72 |
| Posterior insula (L) | 48 | -4.006 | 0.000010 | 70 | -38 | -14 | 6 |
| Inferior temporal gyrus (L) | 20 | -3.238 | 0.000252 | 12 | -42 | -14 | -28 |
| Middle temporal gyrus (R) | 21 | -4.377 | 0.000002 | 232 | 58 | -24 | -8 |
| Inferior temporal gyrus extending to parahippocampal gyrus (L) | 20 | -3.175 | 0.000317 | 32 | -42 | -26 | -18 |
| Postcentral gyrus, medial (R) | 3 | -2.787 | 0.001283 | 15 | 32 | -34 | 50 |
| Postcentral gyrus, medial (L) | 3 | -3.018 | 0.000567 | 18 | -36 | -38 | 56 |
| Fusiform gyrus (L) | 37 | -3.181 | 0.000311 | 88 | -22 | -44 | -18 |
| Middle temporal gyrus (R) |  | -3.156 | 0.000342 | 195 | 48 | -48 | 0 |
| Superior parietal gyrus (R) | 5 | -3.479 | 0.000096 | 195 | 14 | -50 | 70 |
| Middle temporal gyrus (L) | 37 | -4.052 | 0.000008 | 764 | -44 | -68 | 6 |
| Middle temporal gyrus (R) | 37 | -3.774 | 0.000029 | 136 | 42 | -70 | 14 |
| Cerebellum, crus I extending to crus II (L) |  | -2.648 | 0.002050 | 11 | -22 | -82 | -30 |

#### Neural correlates of equitable decisions

In the equitable decision cluster, prosocial decisions (in contrast to selfish decisions) were associated with increased activation in bilateral orbital frontal cortex (OFC), bilateral ventrolateral PFC, bilateral dorsolateral PFC, bilateral medial PFC including rostral ACC, bilateral ventral striatum and caudate, and left occipital cortex. Activation was increased during selfish relative to equitable decisions in left dorsolateral PFC, medial portion of left precentral gyrus, lateral portion of bilateral precentral gyrus, left thalamus, bilateral supramarginal gyrus, left inferior temporal gyrus, bilateral posterior superior temporal sulcus (STS), and bilateral occipital cortex (Figure 3, Table 2).

#### Neural correlates of altruistic decisions

In the altruistic decision cluster, prosocial decisions (in contrast to selfish decisions) were associated with increased activation in several regions including left ventromedial PFC, bilateral ACC and paracingulate gyrus, bilateral pre-SMA, bilateral anterior insula, right ventrolateral PFC, left bilateral dorsolateral PFC, thalamus, right ventral striatum, right precuneus, and bilateral interior parietal gyrus.

We also found increased activation during selfish decision-making, in contrast to altruistic decision-making, in several regions including right dorsolateral PFC, right ventrolateral PFC right putamen, bilateral posterior insula, bilateral precentral gyrus, right middle temporal gyrus (MTG), superior temporal gyrus (STG), right STS left parahippocampal gyrus, right superior parietal gyrus, bilateral middle occipital gyrus, and left cerebellum crus I and II (Figure 3, Table 2).

#### Pairwise meta-analytic map comparisons

Pairwise comparisons confirmed that the meta-analytic maps derived from each cluster of studies were distinct from one another. When comparing activation for cooperative relative to equitable decisions, we found increased activation in regions that included left ventrolateral PFC, left SMA, right caudate, left hippocampus, bilateral thalamus, left VTA, left supramarginal gyrus, and left superior parietal gyrus (Supplemental Figure S2a). We did not find increased activity in any region for equitable relative to cooperative decisions. When comparing activation for equitable relative to altruistic decisions, however, we found increased activation for equitable relative to altruistic decisions in regions that included right ventrolateral PFC, bilateral posterior insula, and right STG (Supplemental Figure S2b).

When comparing activation for altruistic relative to equitable decisions, we found increased activation in regions that included dorsolateral PFC, bilateral pre-SMA, left anterior insula, bilateral caudate, left thalamus, and left hippocampus (Supplemental Figure S2c). Finally, when comparing activation for altruistic relative to cooperative decisions, we found increased activation in regions that included left dorsolateral PFC, left SMA, left MTG, and left angular gyrus (Supplemental Figure S2d). We did not find increased activity in any region for cooperative relative to altruistic decisions. See Supplemental Table S1 for results.

#### Conjunction across meta-analytic maps

Conjunction analyses identified overlapping regions of activation for altruism ∩ equity and cooperation ∩ equity. Overlapping activation for altruism ∩ equity was observed in bilateral dorsolateral PFC (BA9/46; left: [-44, 34, 20], k=19; right: [46, 40, 24], k=24), left ventrolateral PFC (BA10; [-42, 44, -4], k=32), and left visual cortex (BA18, [-20, -96, -8], k=34). Overlapping activity for equity ∩ cooperation was observed in bilateral ventral striatum (left: [-2, 4, -8]; right: [4, 4, -8]; k=4). We did not find overlapping activation for cooperation ∩ altruism nor across all three clusters of studies.

## Data availability

All thresholded and unthresholded meta-analytic activation maps are openly available on the Open Science Framework at https://osf.io/3vu9w/.

## Discussion

Using data from 43 unique fMRI studies that included 25 statistical maps and 18 coordinate tables across 1,423 subjects, we identified 13 features that distinguish prosocial decisions tasks. We used these features to generate a feature-based representation of prosocial decision tasks that classified prosocial decisions into three sub-clusters that we subsequently labeled cooperation, equity, and altruism. That the feature-based structure we generated identifies conceptually and motivationally coherent categories of prosocial decisions was supported by the results of our fMRI meta-analysis, which found evidence suggesting that each category of decision recruits diverging neural systems. The first cluster of decisions (which we labeled as *cooperative*) primarily recruited regions such as dorsal and ventral striatum, VTA, and subgenual ACC. A second cluster of decisions (which we labeled as *equitable*) recruited neural regions such as ventral striatum, dorsolateral PFC, and ventromedial PFC. A third cluster of decisions (which we labeled as *altruistic*) recruited neural regions such as ventral striatum, dorsolateral PFC, ventromedial PFC, pre-SMA, dorsal ACC, and anterior insula.

Our approach demonstrates that the dozens of tasks that have been used to assess the neural correlates of prosocial decisions generally cluster together according to specific shared features. These tasks are adaptations of those used in studies of prosocial decision making outside the scanner and more broadly. We identified key features that distinguish prosocial decisions, including features related to the identity of the beneficiary, the nature of the interaction between agents and beneficiaries, and various outcomes associated with the decision. Then, using an unsupervised graph-based approach, we identified three clusters of tasks that tend to share common core features. For example, cooperative decisions included outcomes that depended on the decisions of others (e.g., “beneficiaries make choices that affect the outcome”) and decisions in conditions of uncertainty. Features shared by equitable decisions included adherence to social norms such as producing equal outcomes and unilateral decisions made by a single agent and that thus resulted in no uncertainty. Features shared by altruistic decisions included outcomes that did not produce any benefit to the deciding agent and unilateral decisions that were made in response to the need or distress of the beneficiaries. Of note, the prosocial decision-making tasks included in this meta-analysis were described by the original authors using at least 10 different terms that did not consistently correspond to the features of the tasks being used—including cooperation, collaboration, reciprocity, trust, equity, fairness, prosocial behavior, interpersonal behavior, charitable behavior, and altruism—reinforcing the value of a clearer and more consistent prosocial decision-making task-space.

Supporting the identified task structure, the clusters yielded by our approach map closely onto the results of a previous behavioral characterization of prosocial paradigms that applied factor analysis to the behavioral outcomes of these paradigms (Böckler et al., 2016). These outcomes included the percentage of prosocial decisions during tasks, the ratio of other-regarding versus self-regarding decisions, average monetary donations, or summary scores of self-reported measures. Using a similar bottom-up, but otherwise completely distinct approach across 329 participants who completed multiple tasks assessing prosocial behavior, the authors identified clusters of prosocial tasks that correspond to those we identified: altruistically motivated prosocial behavior (corresponding to the cluster labeled as *altruistic* decisions), norm-motivated prosocial behavior (corresponding to the cluster labeled as *equitable* decisions), and strategically motivated prosocial behavior (corresponding to the cluster labeled as *cooperative* decisions). They also identified self-reported prosocial behavior (a category not included in our meta-analysis) as comprising a fourth distinct cluster.

Our findings also extend this work by showing that tasks cluster similarly even when completed by different participants across tasks. This suggests task features are crucial in determining the category of a prosocial decision and has implications for comparing results across different tasks, such as in previous meta-analyses. In addition, our findings suggest that even within a type of task, specific features may determine the category of prosocial decision at hand. For example, Dictator Games that offered an option to split available resources equally (50-50) clustered with equitable decisions, whereas Dictator Games with the option to make other prosocial splits clustered with altruistic decisions. The existence of strong norms related to equity may explain why 50-50 is the most common non-selfish split across Dictator Games when this choice is available (Engel, 2011). Future analyses of Dictator Game tasks, particularly those conducted using fMRI, could benefit from considering that there may be something unique about the decision to split resources equally (50%) rather than it simply existing as an option on a parametric continuum between 49% and 51%.

Our approach also yielded several important observations about the neural substrates of prosocial decisions. Notably, all three clusters of prosocial decisions recruited the striatum, but each category of decisions elicited activation in different regions within the striatum. We found that cooperation recruited the left caudate and bilateral ventral striatum, equity recruited right caudate and bilateral ventral striatum, and altruism recruited right ventral striatum. Previous work suggests these differences may be explained by how tasks in the three clusters vary in value and uncertainty. Activity in the striatum has been consistently found to encode action value during learning and decision-making (Daw & Doya, 2006; Guitart-Masip et al., 2014). In some tasks (primarily cooperative and equitable decisions), prosocial decisions increased the agent’s own welfare. In other tasks (primarily altruistic and equitable decisions), prosocial decisions meant forgoing resources. Importantly, in economic games involving simultaneous decisions by multiple agents, cooperative decisions are usually made with some uncertainty about the ultimate outcome. It is possible that the striatal activity observed during cooperative decisions also reflects the uncertainty of decisions, which is recruited during decision-making under risky or uncertain conditions (Farrar et al., 2018; Krain et al., 2006; Lopez-Paniagua & Seger, 2013). The only overlap of striatal activation we identified occurred in a small volume of four voxels during both cooperative and equitable decisions. This conjunction may reflect the fact that cooperative and equitable decisions benefit both agents and beneficiaries, which may be recapitulated in ventral striatal activity.

This suggests the possibility of an additive effect of striatal activity—an interpretation consistent with observations of a parametric effect of striatal activation and reward magnitude for self (Miller et al., 2014). Ventral striatum is also preferentially engaged in response to rewarding social stimuli relative to rewarding nonsocial stimuli, for instance, when participants cooperate with a human partner relative to a computer partner despite identical monetary gains (Rilling et al., 2002, 2004). This also might explain why we did not observe any consistent regions that were more active during selfish decisions than cooperative decisions. Cooperative decisions, which yield outcomes benefiting both the agent and other beneficiaries, are also potentially more rewarding than decisions that only yield self-rewarding outcomes. Cooperative decisions also uniquely recruited activity in bilateral VTA, which projects dopamine to the ventral striatum in response to positive prediction errors and reward cues (D’Ardenne et al., 2008), and activity in which likely reflects the anticipation of both the self- and social-rewards gained from cooperating with others.

Striatal activation to anticipatory reward cues occurs within a larger subjective valuation system, which consistently involves activity in the medial PFC during reward-based decision-making (Bartra et al., 2013) and prosocial decision-making (Bellucci et al., 2020; Cutler & Campbell-Meiklejohn, 2019). As has been previously found (Cutler & Campbell-Meiklejohn, 2019), we observed activation in more anterior portions of the ventromedial PFC (including the rostral ACC) during equitable decisions which produce self-enhancing, norm-based outcomes and activation in more posterior portions during altruistic decisions. These results are consistent with a hypothesized spatial gradient of activation along the medial PFC during prosocial decision-making (Sul et al., 2015), which may integrate information about self and others to encode an overall value during a prosocial decision (Hutcherson et al., 2015).

Activation in both ventral striatum and medial PFC did not overlap between altruistic decisions and cooperative decisions. In contrast to altruistic decisions, which recruited the lingual gyrus portion of left ventromedial PFC, we found activation in bilateral subgenual ACC during tasks that require agents to cooperate with others to achieve a common goal. This suggests that altruistic and cooperative decision represent distinct processes, despite frequent conflation these two categories of decisions in the literature, for example, when altruistic behavior (choosing to benefit others without any self-gain) is labeled “cooperation” (Balliet et al., 2014; Declerck et al., 2013; Gintis, 2014; Peysakhovich et al., 2014; Yang et al., 2019). Because the subgenual ACC supports prosocial learning computations (Christopoulos & King-Casas, 2014; Lockwood et al., 2016) as well as preferences for socially rewarding outcomes (Smith et al., 2010), it may also play a role in updating expectations of others’ actions or the value of others’ outcomes during iterative cooperative decision-making.

In addition to a subjective-valuation sub-system, altruistic decisions seemed to recruit two other distinct sub-systems underlying goal-directed behavior and empathy, respectively. The goal-directed sub-system included regions typically implicated in controlling action and directing goal-directed behaviors, including the lateral PFC (Hoshi & Tanji, 2004; Kaller et al., 2011; Morris et al., 2014). Importantly, activation for equitable and altruistic decision-making overlapped in dorsolateral PFC, involved in modulating subjective value representations (Carlson & Crockett, 2018; Tusche & Hutcherson, 2018) and in making norm-related decisions (Baumgartner et al., 2011; Knoch et al., 2006). Thus, it may play an important role in guiding prosocial action in accordance with abstract, social rules during social decision-making tasks (Bellucci et al., 2020).

The second sub-system comprised regions commonly involved in representing and empathizing with the distress of others, and included the dorsal ACC, pre-SMA, and anterior insula (Ashar et al., 2017; Decety & Lamm, 2006; Lamm et al., 2011). Activation in these regions emerged only during altruistic decisions, consistent with the theory that affective resonance with others’ distress give rise to empathic concern and altruistic motivation, which is a primary motivator of prosocial behavior in the absence of cooperative or equity-maintaining goals (Batson, 2009, 2011; Brethel-Haurwitz et al., 2018; Decety et al., 2016; O’Connell et al., 2019). This finding was likely driven by eight out of the fourteen identified altruistic studies including stimuli that depicted or implied the need or distress of beneficiaries. We also found activation in the precuneus—a key node of the mentalizing system (Koster-Hale et al., 2017)— during altruistic decisions. This region has also been found to be active in response to observing emotional suffering (Immordino-Yang et al., 2009; Masten et al., 2011; Meyer et al., 2013). Although some hypothesize the right TPJ—another core region of the mentalizing system—to be recruited during prosocial decision-making (Chakroff & Young, 2014; Parnamets et al., 2020), we did not find differential activation in this region across prosocial relative to selfish decisions. It is possible that we did not observe differences in mean activation because both selfish and prosocial decisions require the maintenance of others’ beliefs and intentions, whereas studies finding TPJ activation usually contrast decisions pertaining to other people with hypothetical decisions pertaining to imaginary people (or computers) (FeldmanHall et al., 2012) (for further evidence regarding the role of TPJ during domain-general social versus self-specific decision-making, also see: Lockwood et al., 2018, 2020).

## Limitations and Future Directions

These results should be considered in light of some limitations. We could not obtain complete data from a number of potentially relevant studies. At least 69 studies would have been eligible for the analysis if we had been able to retrieve the necessary data. In addition, while the 13 features we identified captured the distinctions across the tasks included in our analysis, they may not be representative of all features that could potentially overlap across tasks. With access to more data, future work could test how clustering algorithms such as the one we employed can generalize to unseen tasks with different combinations of features, or even generate new tasks with unique sets of task features. Another limitation of our approach is that we used a binary coding for our features (1=present, 0=absent), which did not make any prior assumptions about the features (e.g., no assumption that features were parametrically weighted). Therefore, we were unable to test how strongly each feature contributed to the three-cluster solution (since they were all weighted equally).

We identified how tasks were inter-related using a bottom-up feature-based approach (assuming equally important features), and how these inter-relationships give rise to overarching prosocial categories. This is in contrast to the more top-down approaches based on expert-models that have been used to map cognitive constructs like creativity (Kenett et al., 2020), cognitive control (Lenartowicz et al., 2010), or theory of mind (Schurz et al., 2020) onto tasks. Future work could combine these approaches to generate a finer-grained task-space for prosocial decision-making including all possible levels of its cognitive ontology: the categories identified in the present study, finer-detailed sub-categories, task paradigms and their features weighted according to their relative importance, and contrast estimates. In so doing, we could go further in solving discrepancies within the prosocial decision-making literature, such as delineating more specific categories of prosocial decision-making within the identified task-space, which may only reflect the top level of a prosocial decision-making hierarchy. Our data show evidence of such a prosocial decision hierarchy across clusters of studies (see Figure S1). However, future work, with an adequate number of studies on prosocial decisions, will be needed to demonstrate such a hierarchy can be recapitulated using meta-analytic brain maps (see Schurz et al., 2020) (especially with increasing availability of other studies that prompt prosocial decisions). Other categories of decisions likely also exist within this hierarchy. For example, active decisions to forgive (Fourie et al., 2020), norm-enforcing decisions (i.e., social influence on agreements or valuation) (Chang & Sanfey, 2013; Wu et al., 2016; Yang et al., 2019; Zinchenko & Arsalidou, 2018), and third-party altruistic punishment decisions for norm violations (Buckholtz et al., 2008; David et al., 2017; Fehr et al., 2004; Jordan et al., 2016) were not considered in this study because we sought to only examine decisions that directly benefited another person, but may reflect more specific prosocial decisions under the umbrella of the identified categories.

As with many neuroimaging tasks, prosocial decision-making tasks adapted for neuroimaging are tightly controlled, and often designed to minimize variability and maximize the statistical power of detecting effects. However, they are not designed with high ecological validity and may not map onto the contexts of real-world prosocial decisions such as holding a door open, splitting a meal, volunteering, or donating blood or an organ. Instead, they are primarily monetary in nature, repetitive, and may increase behaviors related to social desirability in the laboratory (Richman et al., 1999). Recent work has focused on making neuroimaging paradigms more “naturalistic,” such as viewing or listening to narratives or interacting in real-time with another person in the laboratory (Hasson & Frith, 2016; Redcay & Moraczewski, 2019; Redcay & Schilbach, 2019; Wheatley et al., 2019) or in the real-world (Dikker et al., 2019). Other work has concentrated on characterizing the behavioral and neural features of individuals who engage in extreme forms of real-world prosociality (Brethel-Haurwitz et al., 2018; Marsh et al., 2014; O’Connell et al., 2019; Vekaria et al., 2020). Understanding the neurobiology underlying more ecologically-valid altruistic decisions will be crucial for understanding the broader picture of prosocial decision-making.

Related to this, we were not able to consider how individual differences in phenotypic traits may contribute to the neural activity patterns observed across prosocial decision-making tasks due to the limited number of studies that collect or report consistent data on participant characteristics, yet this remains an open question. Finally, we only considered univariate maps that contrasted prosocial versus selfish decisions (or assessed a parametric increase). Due to the high variability of contrasts across studies, this allowed us to generate consistent neuroimaging contrasts that only indexed activation during the decision phase across studies. This approach is distinct from other meta-analytic work compiling coordinate-based maps across any contrasts in prosocial tasks (Bellucci et al., 2020; Yang et al., 2019), which run the risk of creating dependence across experiment maps that negatively impacts the validity of meta-analytic results (Müller et al., 2018). Ideally, future work would incorporate neural activation maps derived from computational modeling of behavior (Charpentier & O’Doherty, 2018; Lockwood et al., 2020; Lockwood & Klein-Flügge, 2020; Suzuki & O’Doherty, 2020; Tognoli et al., 2017), which holds promise for understanding individual differences in social learning and decision-making (Patzelt et al., 2018). For example, computational modeling of decisions can identify latent subjective states (e.g., mood, anxiety), beliefs about others (e.g., trust, morality), or other subjective biases about agents that are not directly observable from behavior. Thus, mapping these latent parameters onto the task features that give rise to them and their neural representations will be a necessary next step in characterizing a cognitive ontology of prosociality.

## Conclusion

Despite limitations, the present study provides a framework for understanding how prosocial decisions are inter-related and distinct, and can be applied to a variety of experimental task paradigms. Using a bottom-up approach, we identified a feature-based representation of the task-space underlying prosocial decisions. Results revealed that three clusters of prosocial decisions identified this way—cooperative, equitable, and altruistic decisions—recruit neural systems that diverge in ways that shed light on the key motivations and mechanisms that support each category of prosocial decision compared to selfish decisions. These findings clarify some of the existing heterogeneity in how prosociality is conceptualized and generate insight for future research in task paradigm development and the improvement of formal cognitive ontologies.

## Acknowledgements

We would like to thank Peter E. Turkeltaub, Katherine O’Connell, Kathryn Berluti, Adam E. Green, and Joscelin Rocha-Hidalgo for helpful suggestions, discussions, and feedback regarding this project. We are also grateful to Daniel Campbell-Meiklejohn for support in collecting data, Lin Gan, Alexa King, and Kathleen Neill for their help identifying the task features, and all of the researchers who contributed their time and data for our meta-analysis.

## Footnotes

† After conducting this initial search, we conducted an additional PubMed search using the criteria ((“fMRI” OR “neur*”) AND “equity”) and did not identify any studies published prior to 2019 that would have been eligible for the study and were not already included.

