## Supplemental Information for "A feature-based network analysis and fMRI meta-analysis reveal three distinct types of prosocial decisions"

**Table S1.** Pairwise comparisons of meta-analytic activation maps across prosocial categories.

**Cooperation > Equity**

| Region | Brodmann Area | SDM-Z | P | Voxels | MNI-x | MNI-y | MNI-z |
|---|---|---|---|---|---|---|---|
| Inferior frontal gyrus, ventrolateral (L) | 6 | 3.429 | 0.000202 | 91 | -54 | 2 | 14 |
| Supplementary motor area (L) | 6 | 3.297 | 0.000366 | 199 | -4 | -4 | 68 |
| Precentral gyrus (L) | 6 | 3.685 | 0.000061 | 31 | -44 | -8 | 60 |
| Hippocampus (L) | | 3.037 | 0.001108 | 19 | -14 | -12 | -14 |
| Caudate nucleus (R) | | 3.4 | 0.000231 | 27 | 18 | -12 | 24 |
| Thalamus (L, R) | | 3.393 | 0.000239 | 139 | 0 | -22 | 6 |
| Thalamus (L) | | 3.669 | 0.000067 | 152 | -20 | -28 | 8 |
| Cerebellum lobule IV / V extending to fusiform gyrus (R) | 20 | 2.825 | 0.002578 | 10 | 28 | -30 | -26 |
| Ventral tegmental area (L) | | 3.371 | 0.000264 | 144 | -8 | -30 | -26 |
| Supramarginal gyrus (L) | 48 | 2.874 | 0.002123 | 22 | -62 | -32 | 24 |
| Lingual gyrus (R) | 27 | 2.975 | 0.001423 | 16 | 16 | -42 | -6 |
| Supramarginal gyrus (L) | 48 | 2.927 | 0.001731 | 19 | -54 | -42 | 28 |
| Fusiform gyrus (L) | 37 | 3.217 | 0.000518 | 22 | -40 | -48 | -12 |
| Superior parietal gyrus (L) | 5 | 4.157 | 0.000005 | 619 | -18 | -50 | 60 |
| Posterior cingulate cortex, ventral (R) | 23 | 3.421 | 0.000211 | 17 | 30 | -62 | 8 |
| Cuneus (L) | 19 | 3.574 | 0.000105 | 57 | -42 | -62 | 6 |

**Equity > Altruism**

| Region | Brodmann Area | SDM-Z | P | Voxels | MNI-x | MNI-y | MNI-z |
|---|---|---|---|---|---|---|---|
| Inferior frontal gyrus, triangular part (R) | 45 | -2.867 | 0.000067 | 376 | 48 | 36 | 6 |
| Inferior frontal gyrus extending to insula (R) | 6 | -2.726 | 0.000132 | 305 | 48 | -6 | 6 |
| Superior temporal gyrus (R) | 20 | -2.3 | 0.000895 | 37 | 48 | -10 | -14 |
| Posterior insula (L) | 48 | -2.802 | 0.000092 | 70 | -38 | -14 | 6 |
| Superior temporal gyrus (R) | 21 | -2.556 | 0.000292 | 77 | 58 | -22 | -6 |

| | | | | | | | |
|---|---|---|---|---|---|---|---|
| Middle occipital gyrus (R) | 39 | -2.373 | 0.000654 | 29 | 42 | -72 | 26 |
| Cerebellum, crus I (L) | | -2.316 | 0.000832 | 17 | -22 | -80 | -30 |
| Middle occipital gyrus (L) | 19 | -2.215 | 0.001272 | 16 | -34 | -84 | 30 |

**Altruism > Equity**

| Region | Brodmann Area | SDM-Z | P | Voxels | MNI-x | MNI-y | MNI-z |
|---|---|---|---|---|---|---|---|
| Superior frontal gyrus, dorsolateral (L) | 9 | 2.863 | 0.000466 | 17 | -20 | 42 | 44 |
| Anterior cingulate extending to paracingulate gyri (R) | 32 | 2.446 | 0.002611 | 13 | 10 | 36 | 24 |
| Middle cingulate extending to paracingulate gyri (R) | 24 | 2.905 | 0.000385 | 187 | -2 | 22 | 36 |
| Superior frontal gyrus, dorsolateral (L) | 9 | 2.794 | 0.000625 | 32 | -36 | 22 | 32 |
| Anterior insula (L) | 48 | 2.596 | 0.001440 | 24 | -30 | 18 | 10 |
| Supplementary motor area (R) | 8 | 2.435 | 0.002725 | 23 | 12 | 16 | 62 |
| Anterior insula (L) | 48 | 2.63 | 0.001253 | 11 | -34 | 12 | -14 |
| Supplementary motor area (L) | 6 | 2.977 | 0.000280 | 138 | -6 | 6 | 68 |
| Superior middle gyrus, dorsolateral (L) | 6 | 2.664 | 0.001092 | 108 | -48 | 2 | 26 |
| Precentral gyrus (L) | 6 | 2.387 | 0.003274 | 11 | -42 | 0 | 54 |
| Caudate nucleus (R) | | 3.492 | 0.000022 | 98 | 16 | -8 | 22 |
| Thalamus (L) | | 2.573 | 0.001582 | 11 | -6 | -8 | 4 |
| Hippocampus (L) | | 2.556 | 0.001701 | 12 | -18 | -12 | -14 |
| Pons (L) | | 2.959 | 0.000304 | 125 | -10 | -24 | -14 |
| Thalamus (L) | | 3.676 | 0.000009 | 365 | -14 | -30 | 14 |
| Fusiform gyrus (L) | 37 | 2.648 | 0.001169 | 35 | -44 | -46 | -10 |

**Altruism > Cooperation**

| Region | Brodmann Area | SDM-Z | P | Voxels | MNI-x | MNI-y | MNI-z |
|---|---|---|---|---|---|---|---|
| Superior frontal gyrus, dorsolateral (L) | 10 | 2.489 | 0.000941 | 16 | -14 | 62 | 24 |

| | | | | | | | |
|---|---|---|---|---|---|---|---|
| Superior frontal gyrus, dorsolateral (L) | 8 | 2.537 | 0.000803 | 25 | -10 | 38 | 58 |
| Superior frontal gyrus, dorsolateral (L) | 9 | 2.198 | 0.002377 | 21 | -12 | 38 | 46 |
| Supplementary motor area (L) | 8 | 2.322 | 0.001615 | 40 | -10 | 30 | 36 |
| Supplementary motor area (L) | 6, 8 | 2.419 | 0.001180 | 18 | -10 | 24 | 66 |
| Middle temporal gyrus (L) | 21 | 3.869 | 0.000005 | 62 | -66 | -32 | -8 |
| Lateral ventricle (L) | | 2.727 | 0.000421 | 13 | -16 | -32 | 16 |
| Angular gyrus (L) | 39 | 2.574 | 0.000710 | 16 | -40 | -56 | 26 |
| Middle occipital gyrus (L) | 18 | 2.131 | 0.002914 | 21 | -22 | -96 | 0 |

**Figure S1.** Hierarchical dendrogram of the three-cluster solution identified with community detection

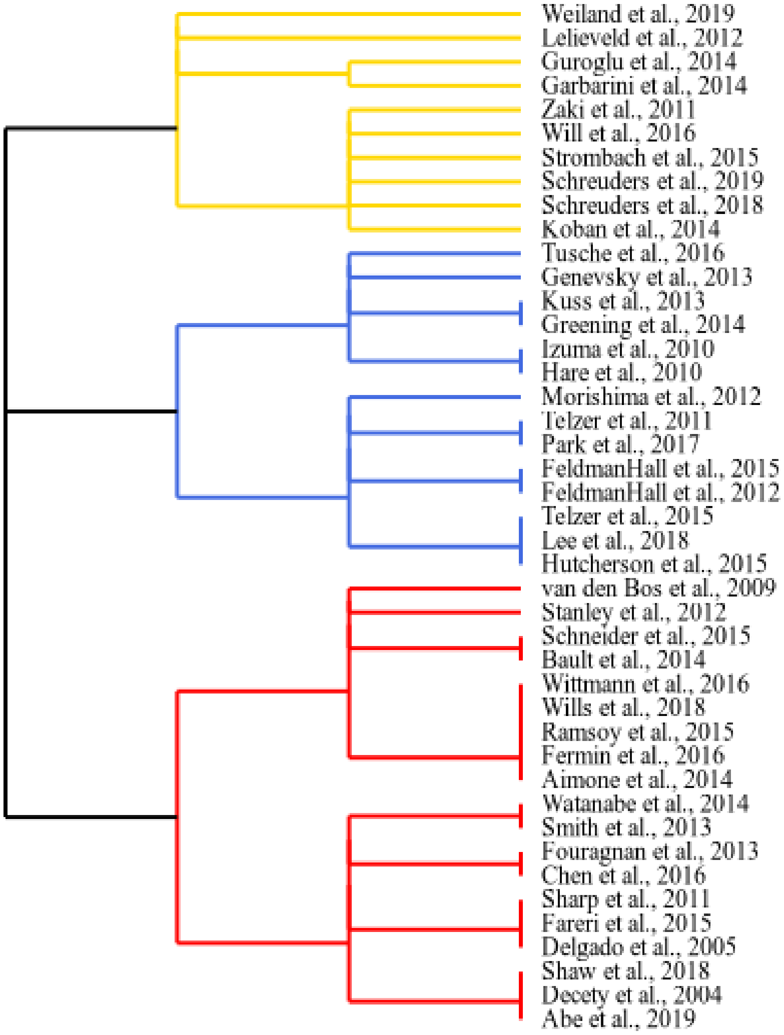


***Note.*** This is a dendrogram depiction of the study network in Figure 2. The hierarchy of relationships were determined using Louvain clustering, which identified three clusters that we labeled cooperative (red), equitable (yellow), and altruistic (blue) decisions based on the task features shared within each cluster.

**Figure S2a.** [Cooperative>Selfish] > [Equitable>Selfish] meta-analytic contrast maps.

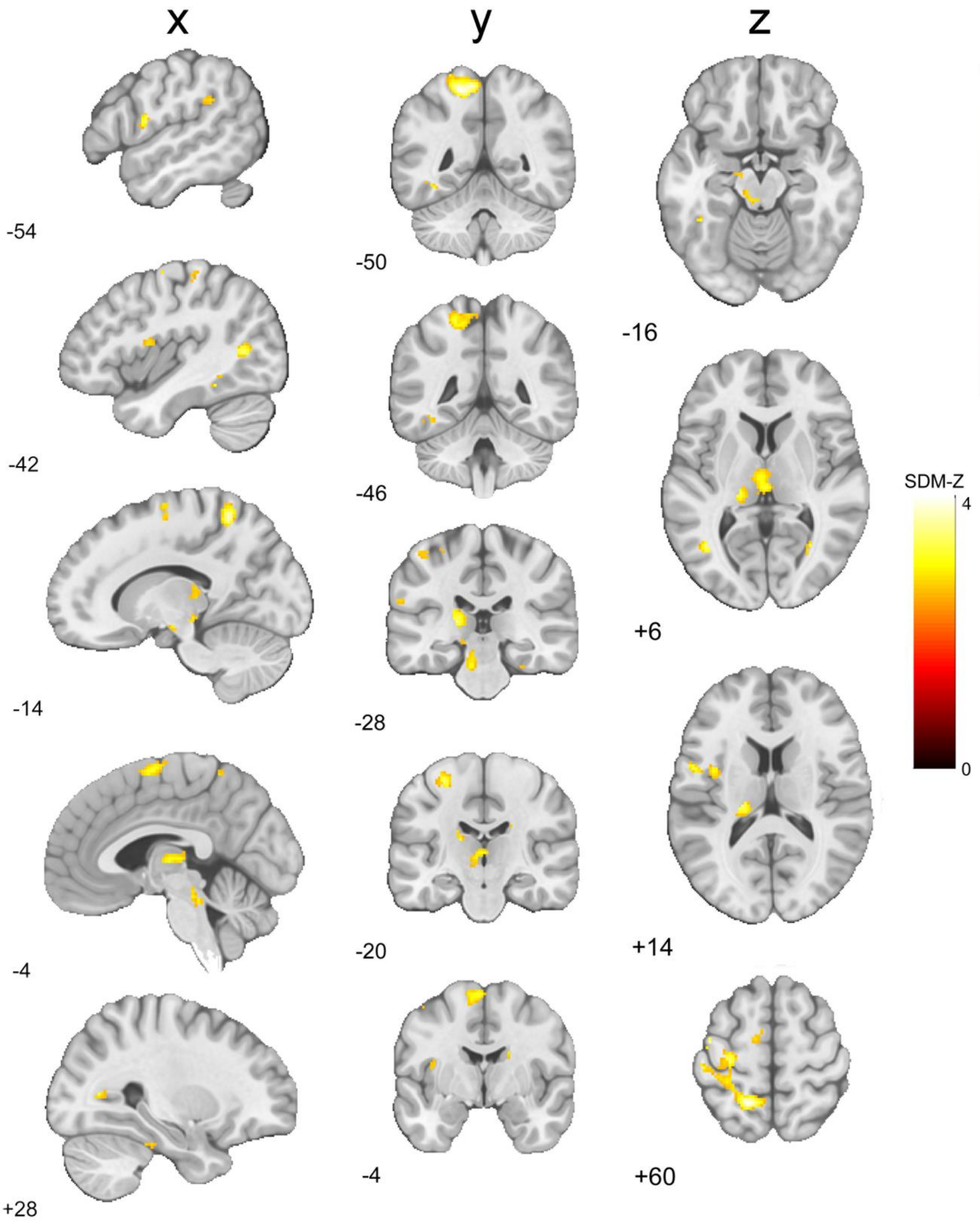

**Figure S2b.** [Equitable>Selfish] > [Altruistic>Selfish] meta-analytic contrast maps.

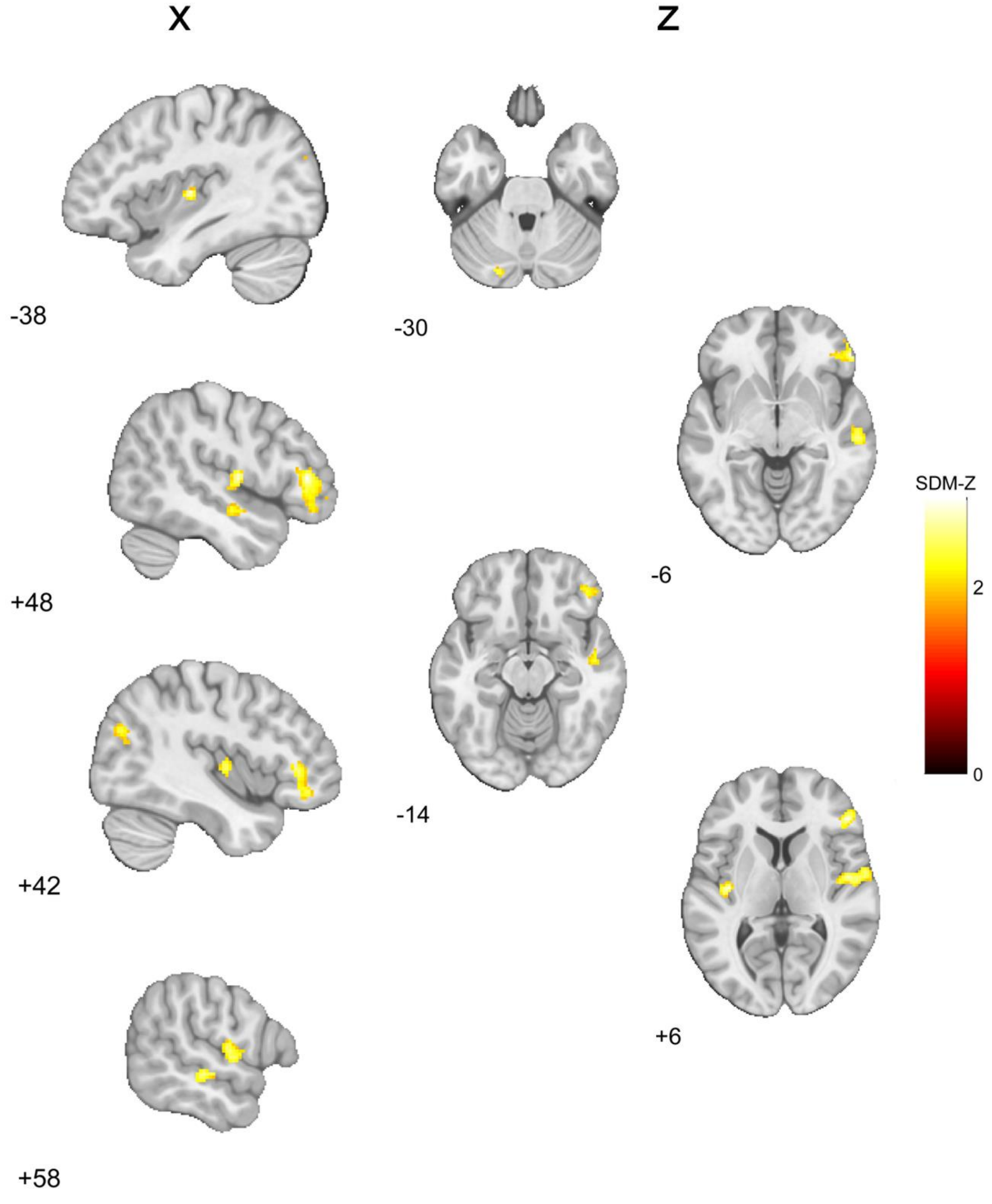

**Figure S2c.** [Altruistic>Selfish] > [Equitable>Selfish] meta-analytic contrast maps.

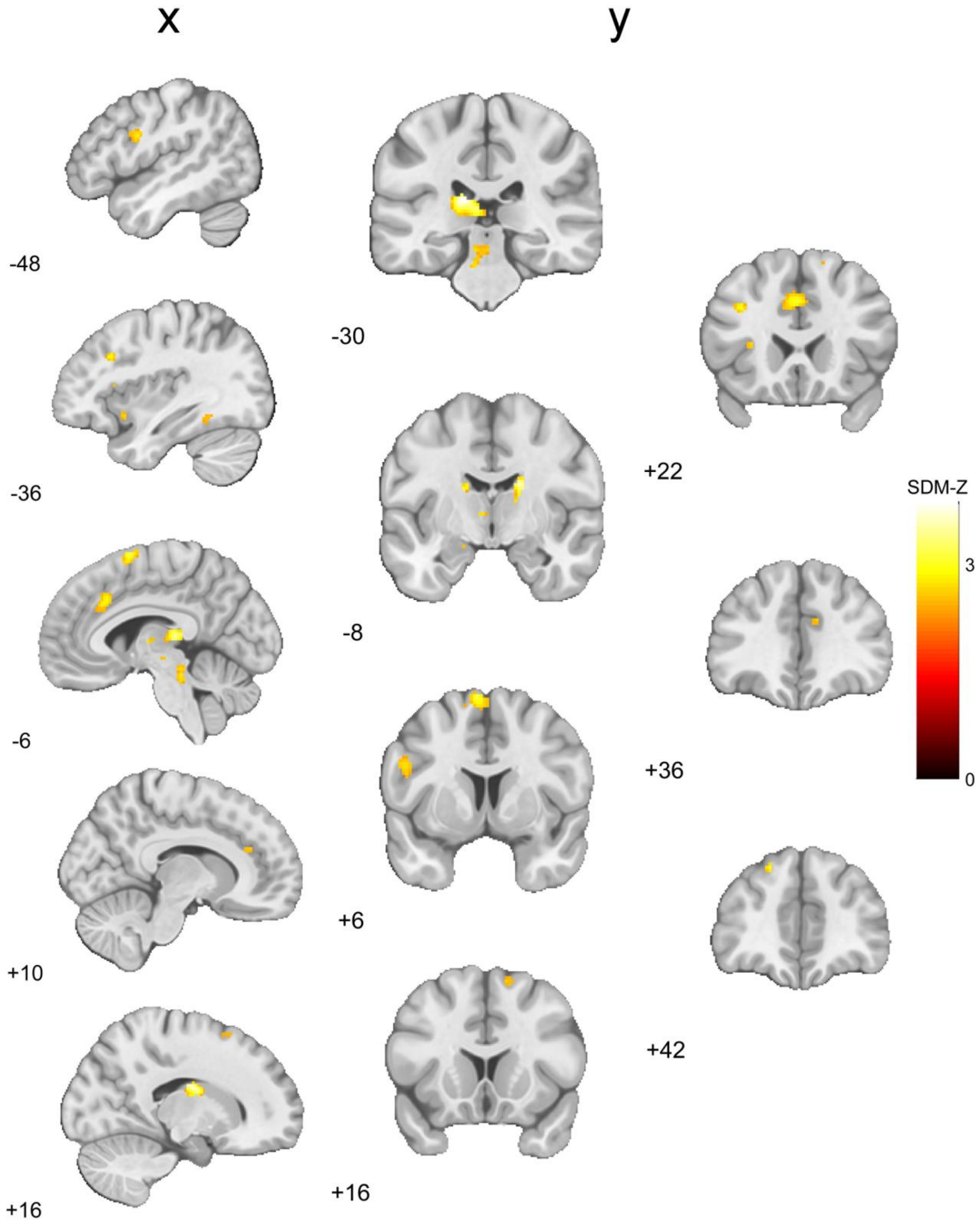

**Figure S2d.** [Altruistic>Selfish] > [Cooperative>Selfish] meta-analytic contrast maps.

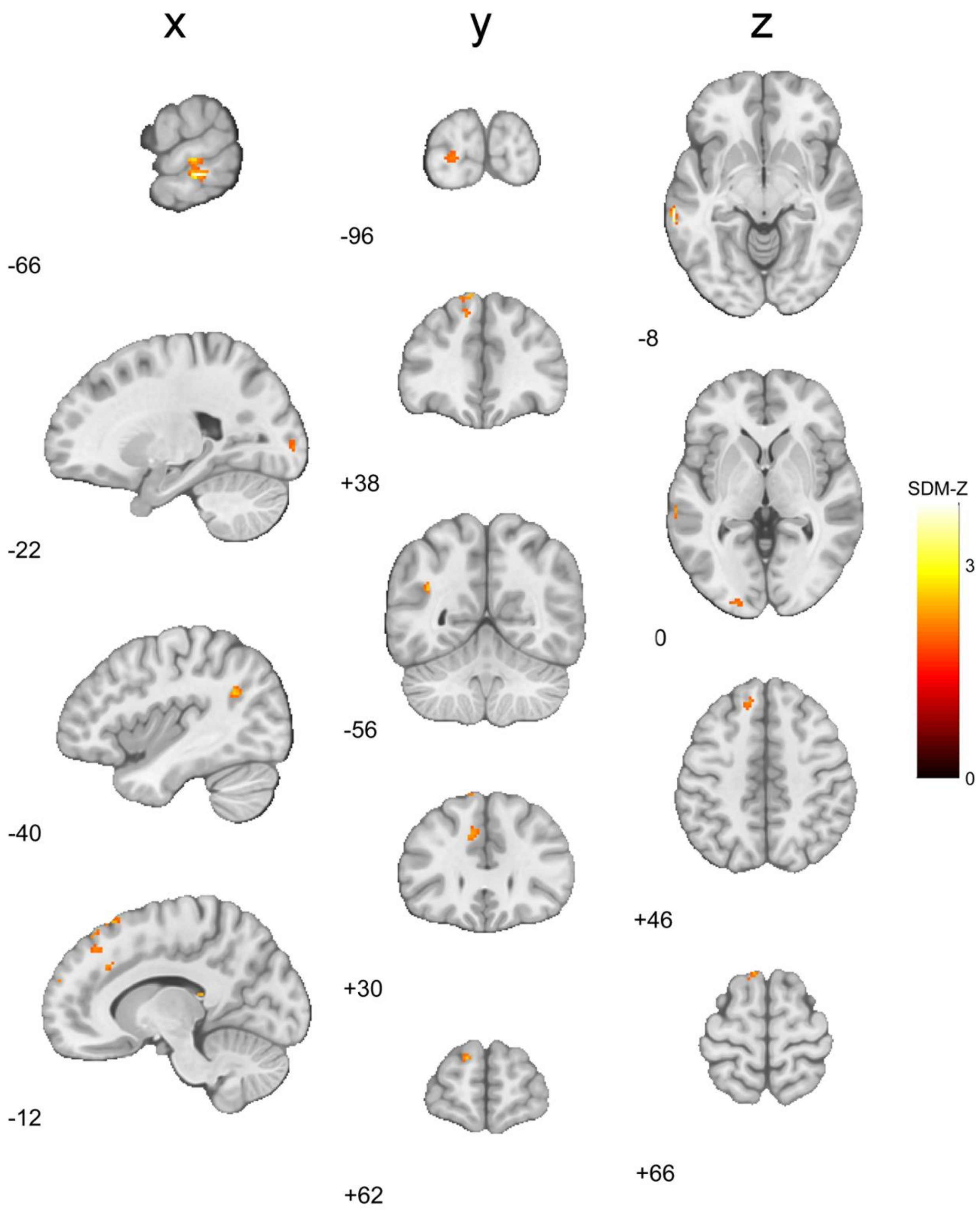
